# Heterogeneous Synaptic Homeostasis: A Novel Mechanism Boosting Information Propagation in the Cortex

**DOI:** 10.1101/2023.12.04.569905

**Authors:** Farhad Razi, Belén Sancristóbal

## Abstract

Perceptual awareness of auditory stimuli decreases from wakefulness to sleep, largely due to reduced cortical responsiveness. During wakefulness, neural responses to external stimuli exhibit a broader spatiotemporal propagation pattern compared to deep sleep. A potential mechanism for this phenomenon is the synaptic upscaling of cortical excitatory connections during wakefulness, as posited by the synaptic homeostasis hypothesis. However, we argue that uniform synaptic upscaling alone cannot fully account for this observation. We propose a novel mechanism suggesting that the upscaling of excitatory connections between different cortical areas exceeds that within individual cortical areas during wakefulness. Our computational results demonstrate that the former promotes the transfer of neural responses and information, whereas the latter has diminishing effects. These findings highlight the necessity of heterogeneous synaptic upscaling and suggest the presence of heterogeneity in receptor expression for neuromodulators involved in synaptic modulation along the dendrite.

## Introduction

During the sleep-wake cycle (SWC), the capacity of the cerebral cortex to transmit neural signals across cortical areas, known as cortical effective connectivity (1), is higher during wakefulness compared to the deep phases of non-rapid eye movement (NREM) sleep (2–4). However, the precise neural mechanisms underlying this enhanced propagation of neural responses remain largely speculative.

Changes in neuronal and synaptic dynamics across the SWC within neural pathways could alter propagation patterns. Experimental evidence indicates that low concentrations of neuromodulators released from the ascending arousal network (AAN) during sleep (5) modulate neuronal dynamics (6, 7) and synaptic strength (8–10). The characteristic oscillating dynamics of neuronal transmembrane voltage, alternating between active (Up) and silent (Down) states during NREM sleep (6, 7), might interrupt communication between cortical regions (11–14). According to this view, evoked Down states following external perturbations in cortical neurons disrupt long-lasting causal interactions among cortical areas during NREM sleep. However, direct experimental evidence supporting this claim remains elusive, and certain empirical observations challenge it as the sole reason for altered cortical effective connectivity during sleep.

For instance, single and multi-unit recordings from the primary auditory cortex (A1) across various species (4, 15–18) have revealed that evoked neural responses to auditory stimuli are comparable across the SWC, despite the aforementioned significant changes in cortical dynamics. It is only in higher-order cortical areas downstream from A1 where evoked neural responses are notably increased during wakefulness compared to NREM sleep (4, 18). This suggests that local changes in neuronal dynamics alone cannot fully account for the differences in response propagation in the cortex, indicating that variations in synaptic strength might also play a significant role.

Regarding changes in synaptic dynamics, the synaptic homeostasis hypothesis (SHY) (9, 10) proposes that synaptic strength in many cortical circuits decreases during sleep to counterbalance the net synaptic upscaling observed during wakefulness. The increase in synaptic strength during wakefulness yields two opposing effects. Firstly, it amplifies stimulus-evoked postsynaptic currents due to the larger projecting axons of excitatory neurons compared to the more localized axons of inhibitory neurons (19–21). Consequently, synaptic upscaling can enhance the amplitude of evoked responses in secondary sensory areas, facilitating the transmission of neural responses across the cortical hierarchy. Secondly, synaptic upscaling enhances spontaneous postsynaptic currents, which are not triggered by external stimuli. *In vitro* studies have shown that an increase in spontaneous synaptic currents, when balanced to avoid overexcitation or overinhibition, decreases the amplitude of evoked responses to external stimuli (22). Therefore, synaptic upscaling simultaneously enhances and diminishes the transmission of neural responses across different cortical areas, depending on the context. This phenomenon underscores the intricate balance of synaptic dynamics and its impact on neural response transmission and processing within the cerebral cortex.

We propose a mechanism that sets up a competition between these opposing effects on evoked neural activities: the *driving effect*, which enhances transmission by amplifying stimulus-evoked postsynaptic currents, and the *pulling effect*, which reduces transmission by increasing spontaneous postsynaptic currents. Uniform synaptic upscaling during wakefulness, without favoring the driving over the pulling effect, may not sufficiently explain the improved propagation of neural responses during wakefulness. The balance between these opposing effects is essential for understanding how re-sponse propagation and information processing occur within the cerebral cortex across different states of consciousness. Hierarchical models of the cortex (23, 24) distinguish between excitatory connections at the circuit level. Generally, inter-excitatory connections link different cortical areas, whereas intra-excitatory connections operate within individual cortical areas. Inter-excitatory connections typically entail a driving effect that facilitates downstream transmission of neural responses, whereas intra-excitatory connections involve modulatory synapses that control local neural activity and lead to a pulling effect.

In this paper, we introduce the *heterogeneous synaptic homeostasis hypothesis* at the circuit level, suggesting that synaptic upscaling should favor inter- over intra-excitatory connections. This approach allows the driving effect, which improves the transmission of neural responses across cortical areas, to prevail over the pulling effect caused by spontaneous postsynaptic currents. The concept of heterogeneous synaptic homeostasis provides a refined perspective on balancing the driving and pulling effects within cortical circuits, emphasizing the significance of the spatial organization of cortical networks that are state–sensitive and facilitate efficient information transmission.

To investigate this hypothesis, we employed a Wilson-Cowan model, which simulates the average firing rate of a cortical column (25). The model replicates dynamics akin to those observed during NREM sleep and wakefulness (26). It has been shown that synaptic upscaling of intra-excitatory connections in a balanced configuration—where augmentation of inhibition counters overexcitation—gradually transitions the spontaneous activity of the model from NREM-like to wakefulness-like dynamics (27). In this study, we examine how adjusting the synaptic upscaling of both intra- and inter-excitatory connections (by factors *β*_intra_ and *β*_inter_, respectively) influences evoked neural responses. Specifically, we study the responses of a single cortical column to stimuli with increasing intensity delivered via inter-excitatory connections. Finally, we explore a scenario where two cortical columns are symmetrically coupled by inter-excitatory connections. One column is perturbed while the other receives stimuli indirectly via inter-excitatory connections from the perturbed to the unperturbed column. We then analyze the effect of varying intra- and inter-synaptic upscaling on the propagation of neural responses between these columns.

Additionally, we establish a framework for quantifying stimulus-relevant information within evoked neural responses and investigate how intra- and inter-synaptic upscaling influence the amount of information that cortical populations convey about a stimulus and its propagation.

## Results

We used a population rate model to simulate the activity of a single cortical column and its interaction with another symmetrically coupled column. By adjusting the strength of excitatory synaptic coupling, we transitioned the model between NREM sleep and wakefulness states. Our analysis concentrated on the effects of synaptic upscaling during these states, particularly focusing on response amplitudes to transient stimuli and the encoding of stimulus intensity in firing responses of pyramidal populations.

### One-cortical-column model

A single cortical column is represented by a model comprising mutually coupled excitatory and inhibitory populations, each receiving independent Gaussian noise inputs (see Fig. 1a).

**Fig. 1.**
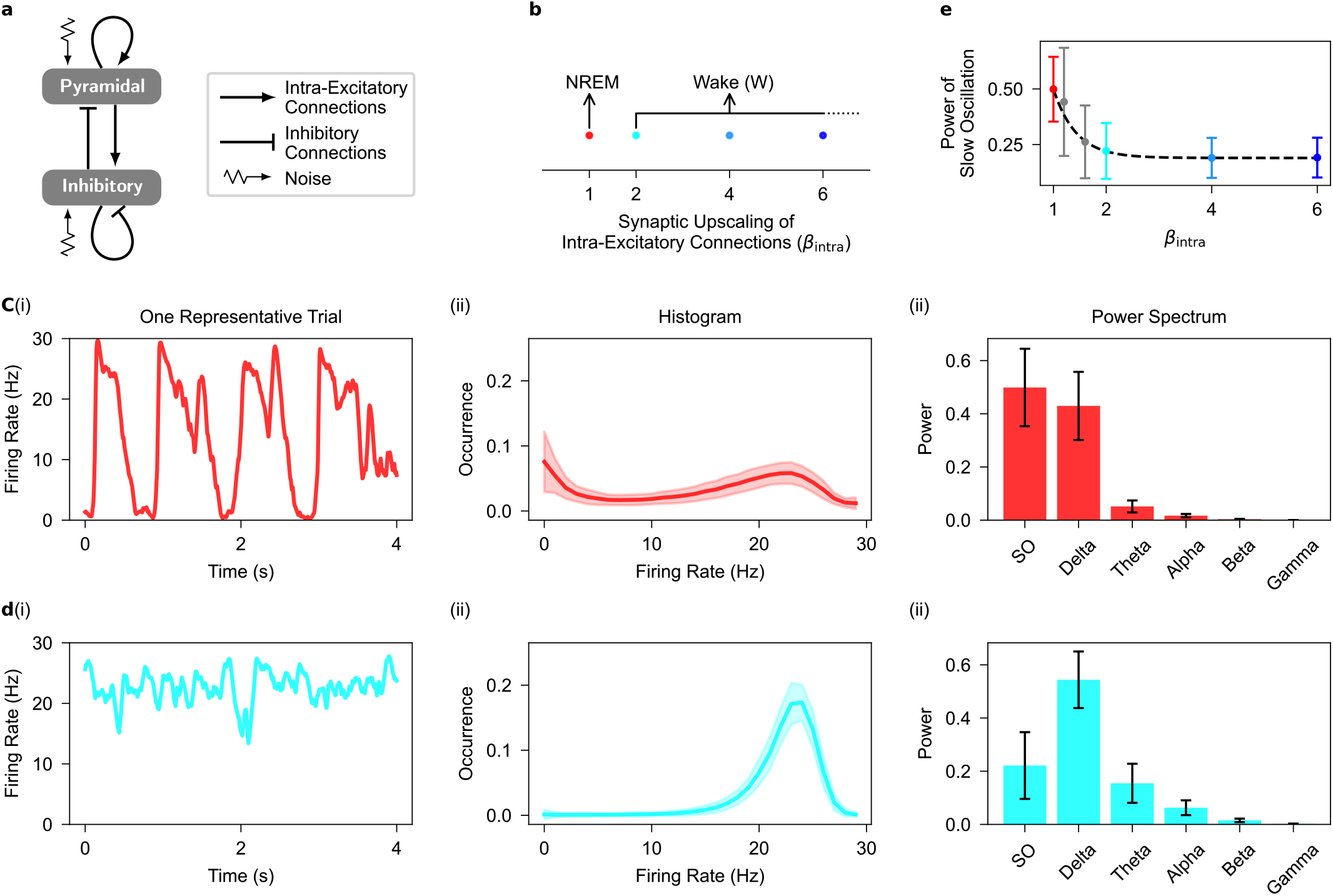
Dynamical features of spontaneous firing activity in the one-cortical-column model. **a**, Diagram of the one-cortical-column model containing one pyramidal and one inhibitory population, where each population receives independent noise. The couplings between pyramidal and inhibitory populations are intra-excitatory and inhibitory connections mediated through, respectively, intra-AMPAergic and GABAergic synapses (see Methods). Refer Table 1 and Table 2 for parameter description and values, respectively. **b**, Parameter space for synaptic upscaling of intra-excitatory connections (*β*_intra_). **c**, Spontaneous firing rate signal for a representative trial (i), the distribution of firing rate signals (ii), and the power spectrum of signals (iii) when there is no intra-synaptic upscaling (*β*_intra_ = 1). The model produces electrophysiological features of NREM sleep when *β*_intra_ = 1. **d**, As in **c**, but for when intra-excitatory connections are upscaled (*β*_intra_ = 2). The model produces electrophysiological features of wakefulness when intra-excitatory connections are upscaled (*β*_intra_ > 1). **e**, The power ratio of slow oscillation (SO) gradually decreases with increasing intra-synaptic upscaling, color coded as in **b**. Shaded area and Error bar correspond to standard deviation over 500 trials. SO, *<*1 Hz; Delta, 1-4 Hz; Theta, 4-7 Hz; Alpha, 7-13 Hz; Beta, 3-30 Hz; Gamma, 30-100 Hz.

#### Electrophysiological patterns

The model parameters (see Table 1 and Table 2) were configured to generate spontaneous firing rates resembling NREM sleep. The parameter for intra-synaptic upscaling was set to 1 (*β*_intra_ = 1; see Fig. 1b), reproducing neural dynamics akin to NREM sleep. These features include high-amplitude fluctuations (see Fig. 1c(i)), a bimodal distribution (see Fig. 1c(ii)), and high power in low-frequency bands (see Fig. 1c(iii)) of the firing rate signals. These features remain robust even when the standard deviation of the noise in the model varies by up to 10% (see Extended Data Fig. 1).

**Table 1.** Parameter description in the one-cortical-column model.

| Symbol | Description |
| --- | --- |
| $Q_k^{\max}$ | Maximal firing rate of population $k$ |
| $\theta_k$ | Firing rate threshold of population $k$ (sigmoid function half activation) |
| $\sigma_k$ | Inverse neural gain of the sigmoid function of population $k$ |
| $\tau_k$ | Membrane time constant of population $k$ |
| $C_m$ | Membrane capacitance in the HH model |
| $\phi_k$ | Gaussian noise on population $k$ |
| $N_{kk'}$ | Mean number of synaptic connections from population $k'$ to population $k$ |
| $\gamma_k$ | Time constant of the postsynaptic response of synapse type $k$, either excitatory or inhibitory |
| $\bar{g}_X$ | Average X-ergic conductance |
| $E_X$ | Reversal potential of the X-ergic current |
| $\bar{g}_{KNa}$ | Average conductance of sodium-dependent potassium channel |
| $E_K$ | Nernst reversal potential of potassium channel |
| $\tau_{Na}$ | Time constant of sodium extrusion |
| $\alpha_{Na}$ | sodium influx through population firing rate |
| $R_{\text{pump}}$ | Strength of the sodium pump |
| $Na_{\text{eq}}$ | Resting state sodium concentration equilibrium |
| $\beta_{\text{intra}}$ | Synaptic-upscaling factor for intra-excitatory connections |
| $\beta_{\text{GABA}}^k$ | Synaptic-upscaling factor for inhibitory connections on population $k$ |

Source: The table is adapted from (26).

**Table 2.** Parameter values in the one-cortical-column model.

| Symbol | Value | Unit |
| --- | --- | --- |
| $Q_p^{\max}$ | 30 | Hz |
| $Q_i^{\max}$ | 60 | Hz |
| $\theta_p, \theta_i$ | -58.5 | mV |
| $\sigma_p$ | 6.7 | mV |
| $\sigma_i$ | 6 | mV |
| $\tau_p, \tau_i$ | 30 | ms |
| $C_m$ | 1 | $\mu\text{F}/\text{cm}^2$ |
| $\phi_{p/i}$ | 1.2 | $\text{ms}^{-1}$ |
| $N_{pp}$ | 144 | — |
| $N_{ip}$ | 36 | — |
| $N_{pi}$ | 160 | — |
| $N_{ii}$ | 40 | — |
| $\gamma_p$ | $70 \cdot 10^{-3}$ | $\text{ms}^{-1}$ |
| $\gamma_i$ | $58.6 \cdot 10^{-3}$ | $\text{ms}^{-1}$ |
| $\bar{g}_{\text{AMPA}}$ | 1 | ms |
| $\bar{g}_{\text{GABA}}$ | 1 | ms |
| $E_{\text{AMPA}}$ | 0 | mV |
| $E_{\text{GABA}}$ | -70 | mV |
| $E_L^p$ | -66 | mV |
| $E_L^i$ | -64 | mV |
| $\bar{g}_{KNa}$ | 1.9 | $\text{mS}/\text{cm}^2$ |
| $E_K$ | -100 | mV |
| $\tau_{Na}$ | 1.7 | ms |
| $\alpha_{Na}$ | 2 | $\text{mM} \cdot \text{ms}$ |
| $R_{\text{pump}}$ | 0.09 | mM |
| $Na_{\text{eq}}$ | 9.5 | mM |
| $\beta_{\text{intra}}$ | 1 | — |
| $\beta_{\text{GABA}}^{p/i}$ | 1 | — |

Source: The table is adapted from (26).

By increasing the strength of intra-synaptic excitatory connections (*β*_intra_ > 1; see Fig. 1b and Table 3), in line with SHY (9, 10), and adjusting inhibitory strengths to prevent overexcitation (see Methods and Table 3), our model gradually transitions from NREM sleep dynamics to wakefulness (see Extended Data Fig. 2). Firing rate signals become low amplitude (see Fig. 1d(i)), show a unimodal distribution (see Fig. 1d(ii)), and exhibit a relative increase in power at highfrequency bands (see Fig. 1d(iii)). Additionally, the model shows a decrease in slow oscillation power (SO, 0.5-1 Hz), a key feature of NREM sleep, with increased intra-synaptic upscaling (see Fig. 1e). These results demonstrate that the model effectively replicates well-established electrophysiological patterns observed during NREM sleep and wakefulness (6, 7, 28–31).

**Table 3.** Parameter values of *β*^*k*^ for intra-synaptic upscalings in wakefulness in the one-cortical-column model.

| | $\beta_{\text{intra}} = 2$ | $\beta_{\text{intra}} = 3$ | $\beta_{\text{intra}} = 4$ |
| --- | --- | --- | --- |
| $\beta_{\text{GABA}}^p$ | 1.961 | 4.724 | 7.488 |
| $\beta_{\text{GABA}}^i$ | 2.165 | 4.65 | 7.134 |

#### Evoked responses to stimuli

To analyze evoked responses, we applied transient stimuli to the cortical column through inter-excitatory connections (see Fig. 2a). These stimuli represent presynaptic firing from an unmodeled upstream pyramidal population and vary in frequency from 10 Hz to 90 Hz in 20 Hz steps (see Methods). Synaptic upscaling was implemented using two distinct scaling factors: *β*_intra_, which modulates intra-synaptic connections within the modeled column, and *β*_inter_, which affects the inter-synaptic connections from the unmodeled upstream pyramidal population to the modeled column. We examined various combinations of *β*_intra_ and *β*_inter_ (see Fig. 2b) to assess their impact on evoked responses in the modeled cortical column.

**Fig. 2.**
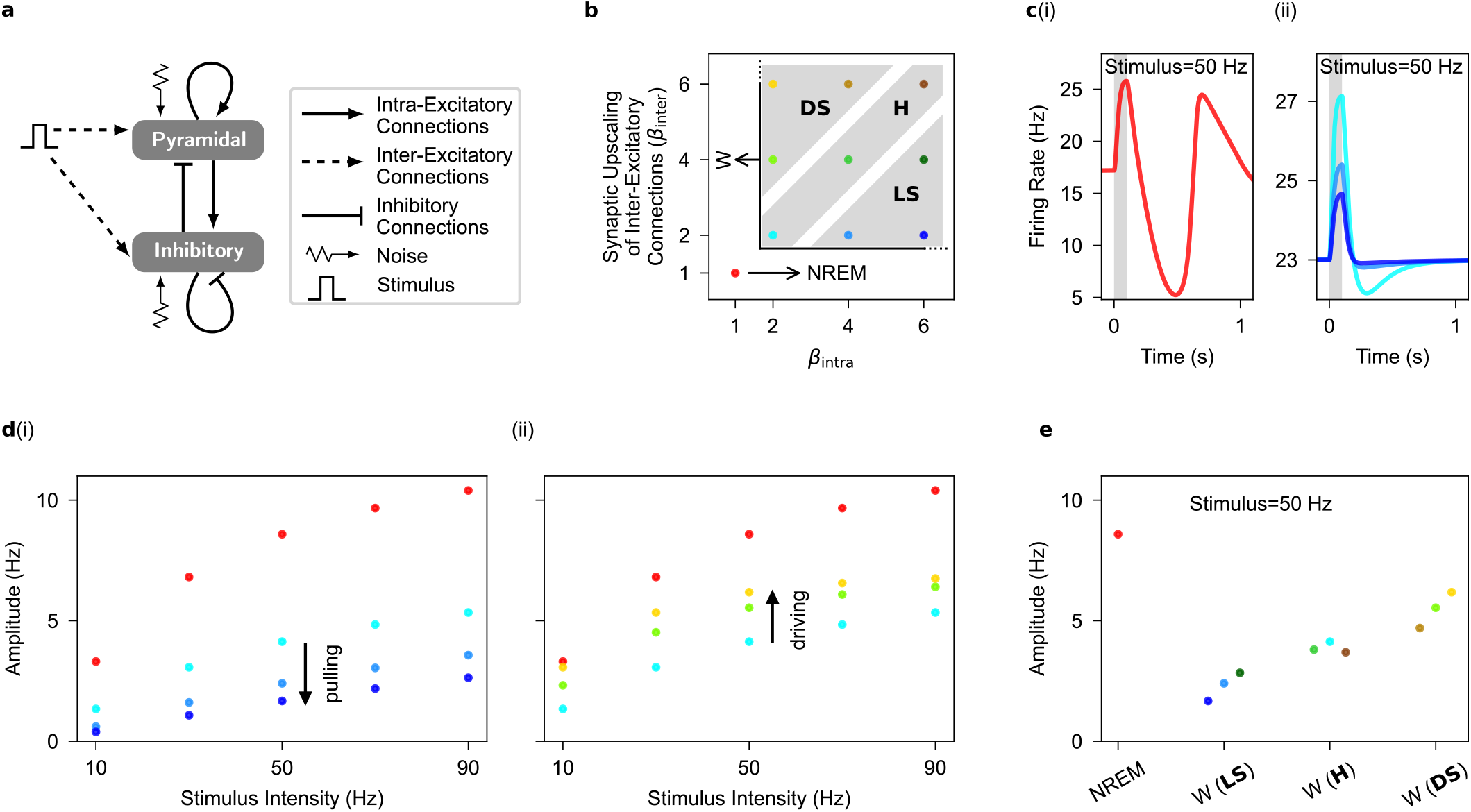
Evoked firing responses to stimuli in the one-cortical-column model. **a**, as in Fig. 1**a**, but for when the model is subjected to stimuli. Stimuli are applied through inter-excitatory connections mediated through inter-AMPAergic synapses. Note that the noise term is set to zero for noise-free evoked response. **b**, as in Fig. 1**b**, but for when intra- and inter-excitatory connections are upscaled in wakefulness. Note that synaptic upscaling in wakefulness can occur under three synaptic upscaling configurations: Local-Selective (LS: *β*_intra_ *> β*_inter_ > 1), Homogeneous (H: *β*_intra_ = *β*_inter_ > 1), and Distance-Selective (DS: *β*_inter_ *> β*_intra_ > 1). **c**, The evoked firing response in NREM sleep (i) and wakefulness (ii), color coded as in **b**, when the stimulus intensity is 50 Hz (the evoked firing responses for other synaptic upscalings in wakefulness are not shown here). Shaded area corresponds to the stimulus duration. **d**, Increasing intra-synaptic upscaling (from *β*_intra_ = 2 to *β*_intra_ = 6) while inter-synaptic upscaling is constant (*β*_inter_ = 2) during wakefulness produces a pulling effect on the amplitude of evoked firing responses (i). On the other hand, increasing inter-synaptic upscaling (from *β*_inter_ = 2 to *β*_inter_ = 6) while intra-synaptic upscaling is constant (*β*_intra_ = 2) during wakefulness produces a driving effect on the amplitude of evoked firing responses (ii). **e**, The amplitude of evoked firing responses increases with increasing values of synaptic upscaling ratio, *β*_inter_ */β*_intra_, during wakefulness. Data points during wakefulness are organized based on increasing values of synaptic upscaling ratio on the x-axis from darkest blue to the lightest yellow. Note that in the homogeneous case, the value of *β*_inter_ */β*_intra_ is equal to one for all three data points. The amplitude of evoked firing responses increases as the synaptic upscaling transitions from local-selective (LS) to distance-selective (DS) upscaling during wakefulness.

In the absence of noise, distinct evoked responses are observed during NREM sleep and wakefulness, consistent with prior experimental observations (16). During NREM sleep, where synaptic upscaling is absent (*β*_intra_ = *β*_inter_ = 1), responses exhibit a wave pattern characterized by an initial surge in firing rate followed by a subsequent decrease below the prestimulus equilibrium, maintaining a steady state well after stimulus offset (see Fig. 2c(i)). In contrast, during wakefulness, synaptic upscaling (*β*_intra_, *β*_inter_ > 1) substantially re-duces the suppression of neuronal firing following activation (see Fig. 2c(ii)).

For all combinations of *β*_intra_ and *β*_inter_, we quantified response amplitudes at stimulus offset. As seen in Fig. 2d, response amplitudes increase with stimulus intensity during both NREM sleep (*β*_intra_ = *β*_inter_ = 1) and wakefulness (*β*_intra_, *β*_inter_ > 1), with NREM sleep generally producing larger amplitudes (red dots in Fig. 2d). Moreover, during wakefulness, keeping *β*_inter_ constant and increasing *β*_intra_ decreases evoked response amplitudes due to the aforementioned pulling effect (e.g., *β*_inter_ = 2 and *β*_intra_ from 2 to 6, shown by light to dark blue dots in Fig. 2d(i)). This pattern holds true across various *β*_inter_ values (e.g., *β*_inter_ = 3 and *β*_intra_ from 2 to 6, shown by light to dark green dots in Fig. 2e). This pulling effect for intra-synaptic upscaling aligns with the reduced amplitude of evoked responses observed in *in vitro* studies as spontaneous postsynaptic currents increase (22).

Conversely, keeping *β*_intra_ constant and increasing *β*_inter_ increases evoked response amplitudes due to the driving effect (e.g., *β*_intra_ = 2 and *β*_inter_ from 2 to 6, shown by the lightest blue, green, and yellow dots in Fig. 2d(ii)). Notably, in the single cortical column architecture, increasing *β*_inter_ exclusively modulates synapses conveying external stimuli, thus not affecting spontaneous postsynaptic currents and preventing a pulling effect.

While both intra- and inter-synaptic upscalings enhance excitatory synaptic currents upon stimulation, the net evoked synaptic current, quantified as |E|−|I|, decreases with increasing *β*_intra_ and increases with increasing *β*_inter_ (see Extended Data Fig. 3a). This highlights the distinct effects of intra- and inter-synaptic upscaling on the evoked synaptic currents during the sleep-waking transition (see Extended Data Fig. 3b). Modulating *β*_intra_ and *β*_inter_ independently allows us to define three distinct synaptic upscaling configurations characterizing the NREM-to-wakefulness transition (see Fig. 2b). These are distinguished by the synaptic upscaling ratio, *β*_inter_*/β*_intra_, as follows:

1. Local-selective upscaling (LS): characterized by *β*_inter_*/β*_intra_ *<* 1.
2. Homogeneous upscaling (H): characterized by *β*_inter_*/β*_intra_ = 1.
3. Distance-selective upscaling (DS): characterized by *β*_inter_*/β*_intra_ > 1.

Presenting findings based on *β*_inter_*/β*_intra_ provides a clearer representation than individual parameters. Firing rate response amplitudes during wakefulness increase with higher synaptic upscaling ratios (see Fig. 2e and Extended Data Fig. 4), reaching maximum values for the DS policy.

#### Stimulus-related information

We quantify stimulus-related information in population firing rates by comparing information detection and differentiation (see Methods and Information Quantification). Information detection quantifies the performance of an optimal classifier in distinguishing evoked responses from spontaneous firing activities. Information differentiation quantifies the performance of an optimal classifier in distinguishing evoked responses elicited by different stimulus intensities from one another.

Information detection increases with stimulus intensity during both NREM sleep and wakefulness (see Fig. 3a). However, increasing *β*_intra_ in the LS policy decreases information detection (light and dark blue dots in Fig. 3a), revealing the pulling effect. Conversely, the driving effect observed with increasing *β*_inter_ in the DS policy during wakefulness enhances information detection beyond NREM levels (light blue and light yellow dots in Fig. 3a, Fig. 3b and Extended Data Fig. 5).

**Fig. 3.**
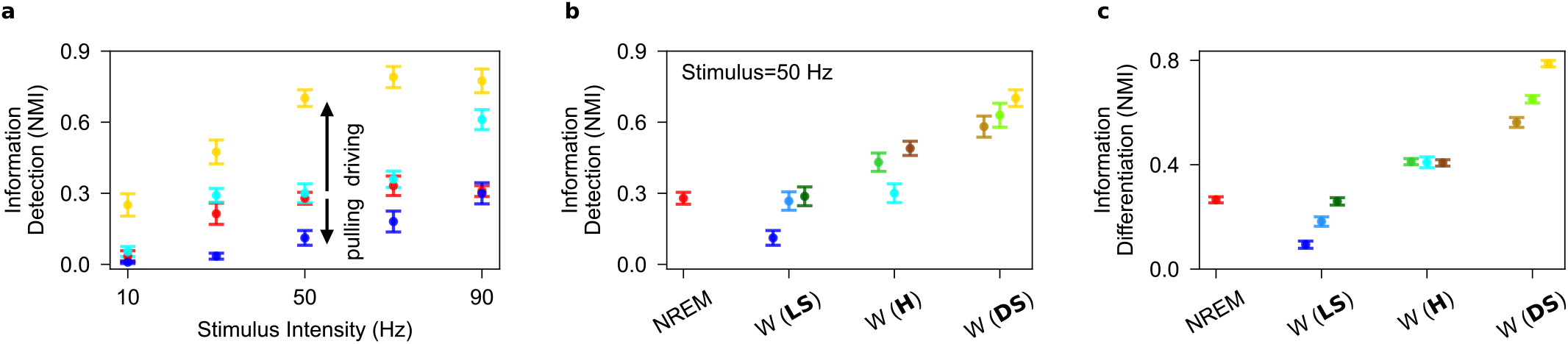
Information content in the evoked firing responses to stimuli in the one-cortical-column model. **a**, Increasing intra-synaptic upscaling while inter-synaptic upscaling is constant (from *β*_intra_ = 2, *β*_inter_ = 2 to *β*_intra_ = 6, *β*_inter_ = 2) during wakefulness produces a pulling effect on information detection. On the other hand, increasing inter-synaptic upscaling while intra-synaptic upscaling is constant (from *β*_intra_ = 2, *β*_inter_ = 2 to *β*_intra_ = 2, *β*_inter_ = 6) during wakefulness produces a driving effect on information detection. **b**, Information detection increases as the synaptic upscaling transitions from local-selective (LS) to distance-selective (DS) upscaling during wakefulness compared to NREM sleep. Note that data during wakefulness is organized based on increasing values of *β*_inter_ */β*_intra_ on the x-axis. **c**, As in **b**, but for information differentiation. Information differentiation increases as the synaptic upscaling transitions from local-selective (LS) to distance-selective (DS) upscaling during wakefulness compared to NREM sleep. Error bar corresponds to 95% confidence interval over 10 performance estimate of the K-means clustering algorithms.

Importantly, the higher trial-averaged amplitudes of evoked responses during NREM sleep compared to wakefulness need to be examined with increased variability across trials. Indeed, the distribution of evoked responses and spontaneous firing activities remain less distinguishable during NREM sleep than wakefulness, compromising information detection in NREM sleep.

Information differentiation during wakefulness exceeds NREM sleep levels under the DS policy (see Fig. 3c). It decreases with increased intra-synaptic upscaling (light and dark blue dots in Fig. 3c), but improves with inter-synaptic upscaling (light blue and light yellow dots in Fig. 3c). This shows consistent encoding of stimulus intensity in firing rates during wakefulness compared to NREM sleep, underscoring the significance of heterogeneous synaptic upscaling favoring inter- over intra-synaptic connections.

### Two-cortical-column model

In this section, we investigate the dynamics of an extended model consisting of two identical cortical columns symmetrically coupled by interexcitatory connections (see Fig. 4a and Table 4).

**Table 4.** Parameter values of connectivity in the two-cortical-column model.

| Symbol | Value | Description |
| --- | --- | --- |
| $N_{pp'}, N_{p'p}$ | 16 | Mean number of synaptic connections from $p$ to $p'$ (and $p'$ to $p$) |
| $N_{ip'}, N_{i'p}$ | 4 | Mean number of synaptic connections from $p$ to $i'$ (and $p'$ to $i$) |

**Fig. 4.**
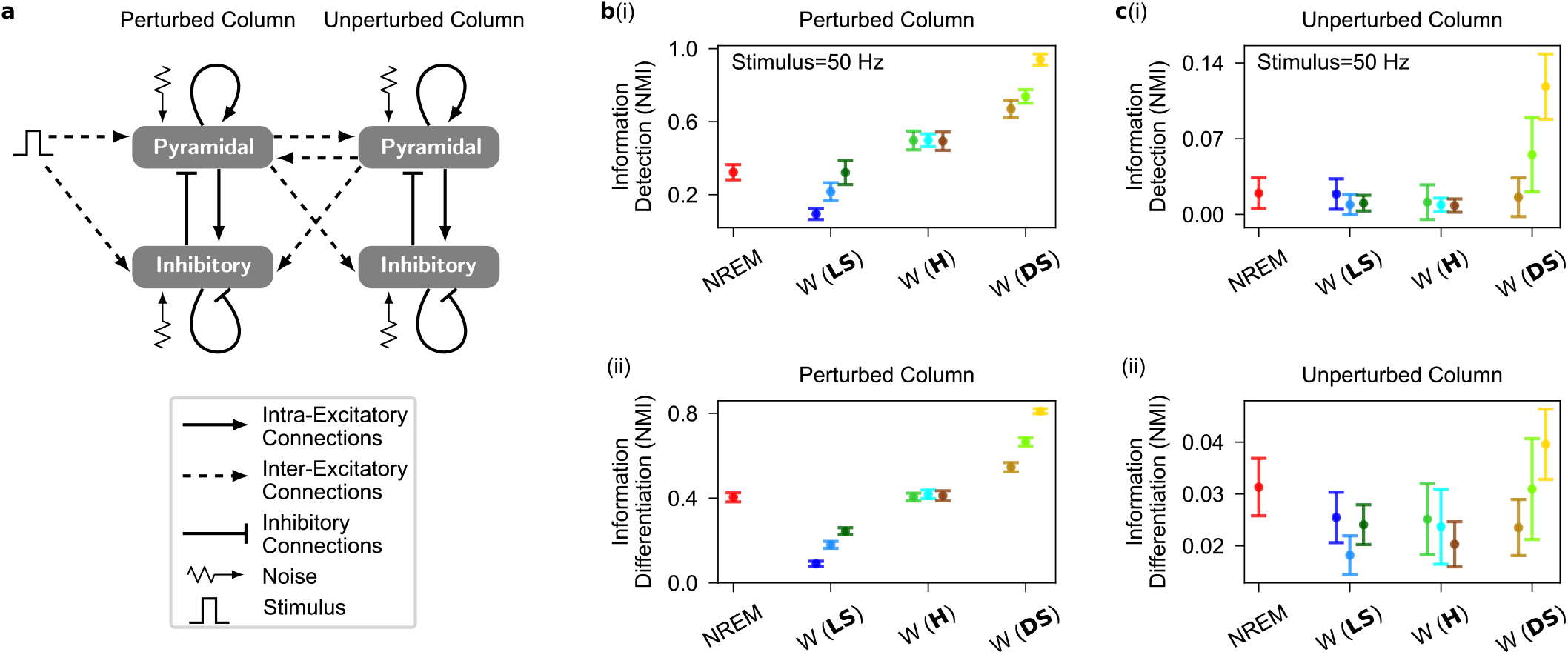
Evoked firing responses to stimuli in the two-cortical-column model. **a**, Diagram of the two-cortical-column model, where each population receives independent noise. The couplings between the two columns are symmetric and are inter-excitatory connections mediated through inter-AMPAergic synapses (see Methods and Table 1, Table 4 and Table 5 for symbol description and parameter values). In the context of spontaneous firing activity, stimulus inensity is set to zero. Note that the noise term is set to zero for noise-free evoked response. **b**, Information detection (i) and differentiation (ii) in the perturbed cortical column increase as the synaptic upscaling transitions from local-selective to distance-selective upscaling during wakefulness compared to NREM sleep. **c**, As in **b**, but for the unperturbed cortical column. Information detection (i) and differentiation (ii) in the unperturbed cortical column increase as the synaptic upscaling transitions from local-selective to distance-selective upscaling during wakefulness compared to NREM sleep. Error bar corresponds to 95% confidence interval over 10 performance estimate of the K-means clustering algorithms.

#### Electrophysiological patterns

The analysis of spontaneous firing activities in the two-cortical-column model replicates the findings of the single-cortical-column model. Increasing *β*_intra_ and *β*_inter_ to mimic synaptic upscaling from sleep to wakefulness (see Table 5) reduces the amplitude of spontaneous fluctuations in pyramidal neuron firing rates (see Extended Data Fig. 6). Notably, the power of the SO band during NREM sleep decreases compared to the single cortical column scenario (see Extended Data Fig. 6a and Fig. 1c), supporting experimental findings that cortical deafferentiation enhances NREM-like dynamics (13).

**Table 5.**
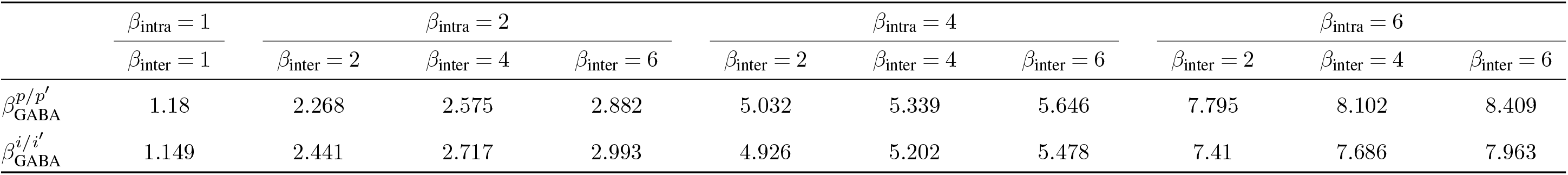
Parameter values of 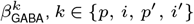, *k* ∈ {*p, i, p′, i′*}, for various synaptic upscalings in the two-cortical-column model.

#### Evoked responses to stimuli

The two-cortical-column architecture serves as a model for exploring downstream information processing from a primary to a secondary cortical sensory area. One column, termed the *perturbed* cortical column, directly receives a stimulus, akin to a primary sensory area receiving direct unimodal thalamic signals. The stimuli modeled in the one-cortical-column model are used here as well. The other column, termed the *unperturbed* cortical column, detects the stimulus solely through presynaptic connections from the perturbed cortical column, mirroring higher-order unimodal sensory areas.

In the absence of noise, the evoked responses of both perturbed and unperturbed populations at stimulus offset reproduce those observed in the single-column model (see Extended Data Fig. 7 and Fig. 2d,e). Increasing *β*_intra_ in the LS policy reduces response amplitudes in both populations (light to dark blue dots in Extended Data Fig. 7a), while increasing *β*_inter_ in the DS policy enhances them (light blue to light yellow dots in Extended Data Fig. 7a). Transitioning from LS to DS upscaling, by increasing the *β*_inter_*/β*_intra_ ratio, boosts response amplitude to external stimuli in both populations (see Extended Data Fig. 7b).

In the two-cortical-column model, both intra- and intersynaptic upscaling boost spontaneous synaptic activity in the unperturbed population, indicating a *pulling* effect now ex-erted by inter-excitatory connections. Nonetheless, intersynaptic upscaling generates an overall driving effect as the net evoked synaptic current increases with increasing intersynaptic upscaling, contrasting with intra-synaptic upscaling (see Extended Data Fig. 7c).

#### Stimulus-related information

Encoding of stimulus intensity in the firing rate of the perturbed population increases from LS to DS upscaling (see Fig. 4b and Extended Data Fig. 8a). DS upscaling is the only policy that increases information beyond NREM sleep levels. In the unperturbed population, both information detection and differentiation greatly surpass NREM sleep levels during DS upscaling (see Fig. 4c and Extended Data Fig. 8b).

These results highlight the necessity for synaptic upscaling across the SWC to be spatially heterogeneous. Specifically, synapses between distinct cortical areas (inter-synapses) must be upscaled more than recurrent synapses within cortical areas (intra-synapses) throughout the SWC. This heterogeneity ensures better stimulus-encoded information during wakefulness compared to NREM sleep across the sensory processing chain.

### Robustness of the Computational Results

To quantify information content, we employed unsupervised machine learning techniques, such as K-means clustering algorithms. Our results remain consistent applying supervised machine learning techniques, such as logistic classification algorithms (see Extended Data Fig. 9a).

Furthermore, our findings are robust across different analytical approaches. Significance tests qualitatively reproduce findings on information detection and differentiation (see Information Quantification and Extended Data Fig. 9b). Moreover, using information theory to compute the mutual information (MI) between the distribution of evoked responses at the stimulus offset and the distribution of stimuli reveals that MI increases as synaptic upscaling transitions from LS to DS upscaling policy (see Extended Data Fig. 9c).

## Discussion

Substantial differences exist in the discharge pattern of the AAN in the brainstem across various states of consciousness (32, 33), resulting in alterations in neuromodulator concentrations throughout the brain (5). These molecular changes impact neuronal (6, 7) and synaptic (8–10) dynamics, potentially altering the cortex’s ability to efficiently transmit neural signals (1–4). Nevertheless, the direct causal relationships between these biological layers remain unclear.

This research investigates the relationship between the synaptic upscaling of excitatory connections (8–10) and enhanced cortical effective connectivity (2–4) during the transition from NREM sleep to wakefulness. Through computational modeling, we offer insights into how synaptic upscaling of excitatory connections not only induces dynamic changes in the electrical activity of the neural networks but also alters information propagation across these networks. Our results show that a spatially broader propagation of information and neural responses occurs during wakefulness compared to NREM sleep, provided that synaptic upscaling between distinct networks surpasses that of local and recurrent connections.

Our result aligns with several previously published computational studies. Firstly, the strengthening of inter-areal excitatory connections in a balanced configuration has been shown to enhance signal transmission in a network model of the macaque cortex (34). Secondly, this outcome aligns with findings that rare long-range connections are necessary for information processing (35).

Our study concentrates on the interaction between two cortical columns, excluding whole-brain interactions. A more detailed model including various cortical and subcortical struc-tures might offer additional insights into the spatial distribution of synaptic scaling. In our simplified model, the connections between the two cortical columns are excitatory and symmetrical, potentially not capturing the full complexity of structural connectivity across all cortical regions. Future work could benefit from distinguishing between feedforward and feedback excitatory connections to better understand how distinct modulations of synaptic upscaling affect propagation of information and neural response. Nonetheless, these would not invalidate the core concept of the DS synaptic upscaling policy as our conclusions hold true even when limited to a single cortical column, showing that DS synaptic upscaling improves stimulus-induced information encoding. Moreover, our neural mass model presupposes that neural communication is based on rate coding. Future investigations could explore spiking-based models since our hypothesis is not conditioned by the coding scheme (temporal or rate based).

A significant contribution of our study is the development of a framework for measuring information content within neural signals. Most studies in sleep research have not explicitly evaluated information content and are primarily based on the amplitude of evoked neural signals. Our work addresses this gap by showing that neither information encoding nor its propagation are enhanced in wakefulness over NREM sleep, except when inter-synaptic upscalings surpass intra-synaptic upscalings. The robustness of our findings is supported by employing a variety of analytical methods, including unsupervised and supervised machine learning techniques, alongside statistical tests and information theory.

Recent studies highlight the significance of analyzing informational content. Research using Neuropixels probes in mice shows that burst firing in thalamic neurons and ampli-tude of cortical responses to electrical stimulation are highest during quiet wakefulness and lowest during anesthesia (36). Since thalamic neurons tend to fire in bursts during NREM sleep (37–39), we might expect similar high activity levels in thalamic relay neurons during NREM sleep as during quiet wakefulness, despite reduced cortical response amplitudes. Our research suggests that distinguishing between information detection and differentiation in stimulus-evoked activity could explain how cortico-thalamo-cortical connections adjust information transfer during quiet wakefulness and NREM sleep.

Wakefulness is associated with consciousness and the capacity to respond to environmental stimuli, whereas sleep diminishes sensory perception (40–42). Human EEG studies show the sleeping brain can perform basic auditory tasks, although higher cognitive functions are compromised. For instance, cognitive response to the subjects’ own name during sleep is similar to that observed during wakefulness (43), whereas motor preparation in response to auditory stimuli are attenuated during NREM sleep relative to wakefulness (41). Moreover, sensory encoding of intelligible stories attenuates during NREM sleep compared to wakefulness, despite comparable encoding of unintelligible stories (42). These results point toward the diminished capacity of the brain to process sensory information with a higher cognitive demand during NREM sleep. Our findings suggest that heterogeneous synaptic upscaling from sleep to wakefulness enhances information detection and differentiation across a broader region of the cortex, allowing for more complex cognitive computations.

Our findings suggest reevaluating the SHY with a focus on circuit-level heterogeneity. Although our results need further empirical support, evidence exists for cellular-level heterogeneity in synaptic upscaling, particularly between perforated and non-perforated synapses. Perforated synapses are larger with discontinuous post-synaptic densities (PSDs), while non-perforated ones are smaller with continuous PSDs (44). Perforated synapses in the mouse cerebral cortex expand their axon-spine contact area upon waking, unlike non-perforated synapses (44). This structural difference highlights a selective approach to synaptic homeostasis at the cellular level. Further experiments are required to explore if cellular-level synaptic homeostasis heterogeneity aligns with our circuit-level hypothesis.

Heterogeneous synaptic upscaling increases the recruitment of a wider network of neural populations across the cortical hierarchy during wakefulness compared to sleep, which is necessary for the emergence of various collective computations within networks of interconnected neurons (45). Yet, the mechanism behind heterogeneous synaptic homeostasis remains unclear. We offer a speculative explanation: intrasynaptic and inter-synaptic connections lie on different dendritic segments, each with a distinct receptor density for neuromodulators secreted by the AAN. Thus, the heterogeneity in receptor expressions results in the heterogeneous synaptic homeostasis during the SWC.

In a wider perspective, heterogeneity seems to be the norm rather than the exception within the brain. For instance, neural firing in pyramidal cells differs based on their target destinations, indicating heterogeneity within traditional pyramidal cell types (46). Moreover, the developmental and regional distribution of N-methyl-D-aspartate (NMDA) receptors have been observed to be heterogeneous (47).

Research indicates that neural heterogeneity plays a functional role in the brain. It improves information transfer in spiking neural networks (48), enhances coding efficiency in predictive coding models (49), acts as a homeostatic mechanism preventing seizures (50), and supports stable learning in recurrent neural networks (51). Our findings contribute to this field by showing that the increased cortical effective connectivity observed during wakefulness relative to NREM sleep may arise from heterogeneous synaptic homeostasis.

## Methods

Our study adapts the Wilson–Cowan model (25) to simulate cortical column activity during both NREM sleep and wakefulness (26, 27).

### One-Cortical-Column Model

A cortical column is represented by interacting pyramidal (*p*) and inhibitory (*i*) neuronal populations (see Fig. 1a). The temporal evolution of average membrane potentials (*V*_*p/i*_) for each population was modeled using a conductance-based approach (52):

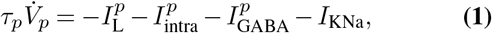

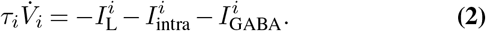

In these equations, *τ*_*k*_ denotes the membrane time constant, while 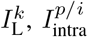 and 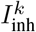 represent the leak, intra-excitatory, and inhibitory currents, respectively, for population *k* (*k* ∈{*p, i*}) as follows:

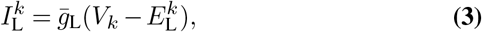

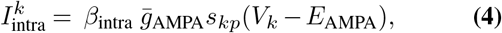

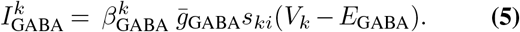

For detailed parameter descriptions and values, refer to Table 1 and Table 2. In brief, 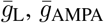, and 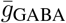 represent average conductances for leak, AMPA-ergic, and GABA-ergic channels, respectively. 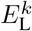, *E*_AMPA_, and *E*_GABA_ denote their corresponding reversal potentials. *β*_intra_ represents intra-excitatory synaptic upscaling, while 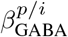 adjusts inhibitory synaptic strength to counterbalance increased excitation due to the synaptic upscaling. This procedure is in line with inhibitory synaptic plasticity of neighboring excitatory synaptic plasticity (53, 54). *s*_*kk*_ ′ represents the synaptic response in population *k* due to presynaptic activity from population *k*′. It is formulated as the convolution of presynaptic firing rate *Q*_*k*_′ with the average synaptic response, characterized by an alpha function with time constant *γ*_*k*_′ (55), following the second-order differential equation:

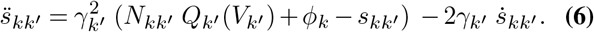

*N*_*kk*_′ represents the connectivity from population *k*′ to *k. ϕ*_*k*_ represents noise and is applied via intra-excitatory connections, simulated independently for each cortical population as a Gaussian process with zero autocorrelation time, zero mean, and 1.2 ms^−1^ standard deviation. Firing rates of population *k* is modeled using a sigmoid function of the average membrane potential (25) as:

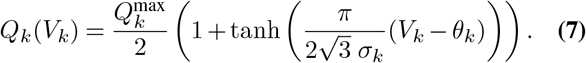

Where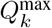, *θ*_*k*_, and *σ*_*k*_ represent the maximum firing rate, threshold, and inverse neural gain of population *k*, respectively.

An activity-dependent potassium current, *I*_KNa_, is included in pyramidal populations to produce NREM-like dynamics (28, 29, 31):

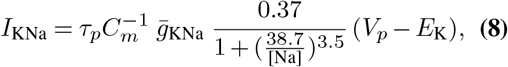

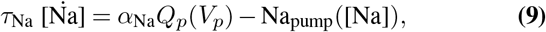

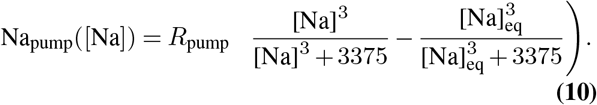

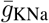 and *E*_K_ are the average conductance and reversal potential of the activity-dependent potassium channel. *C*_*m*_ is membrane capacitance. [Na] represents sodium concentration, with *τ*_Na_ as its extrusion time constant. *α*_Na_ denotes sodium influx due to firing, while Na_pump_([Na]) represents sodium extrusion through pumps with strength *R*_pump_. [Na]_eq_ is the equilibrium sodium concentration.

The model exhibits NREM-like dynamics when *β*_intra_ = 1 (see Table 2 for parameter values). To simulate wakefulness, we increased intra-excitatory synaptic strength (*β*_intra_ > 1), aligning with SHY (9, 10). We balanced this upscaling by en-hancing inhibitory synapses (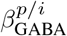; see Table 3) to prevent overexcitation (53, 54). This approach maintains steady-state average membrane potentials for both pyramidal (*V*_*p*_) and inhibitory (*V*_*i*_) populations, preserving excitation-inhibition balance across different synaptic upscaling factors (see Computational pipeline).

Stimuli are delivered via inter-excitatory connections, representing presynaptic firing of an unmodeled pyramidal population, 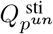. This external stimulation (see Fig. 2a) induces an excitatory synaptic current 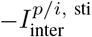 in both pyramidal and inhibitory populations that represents the stimulus-induced excitatory current and follows:

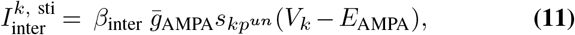

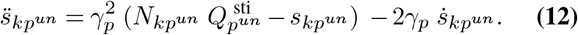

*β*_inter_ is the synaptic upscaling factor for the inter-excitatory connections. 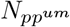 and 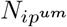 represent the mean number of synaptic connections (16 and 4, respectively).

### Two-Cortical-Column Model

We extended the model to two bidirectionally connected cortical columns (see Fig. 4a), implementing symmetric excitatory connectivity between them (see Table 4), consistent with previous studies (19, 21, 56–59). The average membrane potentials in this expanded model evolve according to:

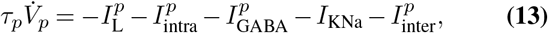

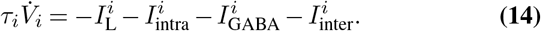

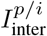 represents the inter-column excitatory currents in our two-column model as:

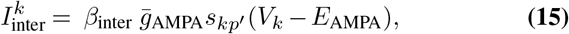

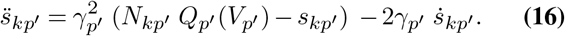

*p*′ represents the pyramidal population in the second cortical column. The second column’s dynamics mirror the first, with population indices swapped (*p* with *p*′ and *i* with *i*′).

As before, synaptic upscaling maintains E/I balance between intra- and inter-synaptic upscalings (see Table 5). An external stimulus applied via inter-excitatory synapses (see Fig. 4a) directly affects only one cortical column, referred to as the *perturbed column*, inducing an excitatory current 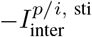, where 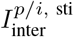 follows Equation 11. The second column, referred to as the *unperturbed column*, receives the stimulus indirectly through inter-excitatory connections from the perturbed column.

### Computational pipeline

We implemented our simulations in Python, employing a stochastic Heun method (60) with a temporal resolution of 0.1 ms. The complete codebase is publicly accessible on GitHub (61). Each trial was simulated independently, with all variables initialized using random values drawn from a uniform distribution. To ensure steady-state dynamics, we discarded the initial 4 seconds of each simulation to eliminate transients. Our analysis focused on the subsequent 4-second period of stabilized activity.

To achieve balanced synaptic upscaling, we conducted 500 independent trials across the parameter space representing NREM sleep (see Table 2). From these simulations, we extracted the peak membrane potentials of pyramidal and inhibitory populations during Up states, identified as the active modes in the bimodal distribution of spontaneous firing activities. Down states, conversely, corresponded to the silent modes. We then calibrated the strength of inhibitory synapses (i.e.,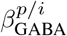; see Table 3) for synaptic upscalings in wakefulness (*β*_intra_) to ensure that the steady-state average membrane potentials of both pyramidal (*V*_*p*_) and inhibitory (*V*_*i*_) populations matched their respective Up state values during NREM sleep. This approach maintains neuronal excitability across sleep-wake transitions and aligns with experimental observations that cortical neural activity during Up states mirrors those during wakefulness (7, 62). This approach is also repeated for the two-cortical-column model (see Table 5).

Our study encompassed both stochastic and deterministic simulations. For stochastic simulations, we introduced Gaussian noise and performed 500 independent trials for each brain state, including NREM sleep and various synaptic upscalings (*β*_intra_, *β*_inter_ > 1) in wakefulness. Additionally, we conducted deterministic simulations without the Gaussian noise to evaluate the model’s response amplitude to external stimuli under controlled conditions.

### Data Analysis

All data analyses were conducted offline using Python. Our analysis primarily focused on the firing rate signals of the pyramidal populations.

#### Analysis of Spontaneous Electrophysiological Patterns

The dynamic characteristics of spontaneous activities were assessed across 500 independent trials for each brain state (see Fig. 1c,d). To quantify the amplitude variability of these spontaneous activities, we computed normalized distributions of firing rate signals for each distinct brain state (see Fig. 1c(ii)). Additionally, we employed Welch’s method (63) to generate spectrograms of the firing rate signals and to characterize the frequency content of spontaneous activities in each brain state (see Fig. 1c(iii)).

#### Analysis of Evoked Responses to Stimuli

The amplitude of evoked firing responses was extracted at the stimulus offset from the deterministic simulations by subtracting the prestimulus values (see Fig. 2d). We also calculated synaptic excitation (E) and inhibition (I) on pyramidal populations across different brain states (see Extended Data Fig. 3). It is important to note that in stochastic simulations, as the number of trials approaches infinity, the amplitudes of evoked responses and E/I values converge to those observed in deterministic simulations.

#### Analysis of Stimulus-Related Information

We developed a comprehensive framework to assess stimulus-related information in stochastic evoked firing responses (see Information Quantification). Our approach is grounded in the neuronal perspective of information as “a difference that makes a difference” (64). To quantify this information in neural firing responses, we introduce two novel measures: *information detection* and *information differentiation*.

*Information detection* quantifies the statistical distinction between stimulus-evoked neural firing responses and spontaneous activities. This measure evaluates whether observed firing patterns can be reliably attributed to stimulus presentation. While crucial for perception, information detection alone does not ensure rich encoding of external stimuli in neural responses. *Information differentiation*, on the other hand, assesses the statistical dissimilarity among neural firing responses to various stimuli. This measure determines whether distinct firing patterns can be reliably associated with specific stimuli, indicating the neural system’s capacity to discriminate between different stimuli.

High levels of information detection and differentiation facilitate precise decoding of stimulus features from neural firing patterns by an ideal observer possessing prior stimulus knowledge. To quantify the information content within stochastic evoked responses, we employ a diverse set of analytical approaches, including machine learning algorithms, statistical significance tests, and information-theoretic methods. For researchers interested in replicating or extending our analysis, we have made our custom Python module for information quantification, *iQuanta*, publicly available on GitHub (65).

#### Unsupervised Machine Learning framework

Our unsupervised machine learning framework utilized the K-means clustering algorithm. To quantify information detection, we applied K-means clustering independently for each stimulus intensity in each brain state, aiming to distinguish stochastic evoked firing responses to a specific stimulus intensity at stimulus offset from spontaneous firing activities. We assessed the clustering algorithm’s efficacy using Normalized Mutual Information (NMI), an external validation metric (66). NMI quantifies the correspondence between predicted cluster labels and true labels, with values ranging from 0 (indicating random clustering) to 1 (denoting perfect clustering).

To evaluate our algorithm’s performance, we implemented 10-fold cross-validation (67). We report the average performance across these 10 folds, accompanied by the 95% confidence interval, for information detection (see Fig. 3a). For assessing information differentiation, we again employed K-means clustering method, this time to distinguish between stochastic evoked firing responses to various stimulus intensities at stimuli offset. This process was conducted independently for each brain state. As with information detection, we quantified performance using NMI, reporting the average across 10 cross-validation folds along with the 95% confidence interval (see Fig. 3c).

For a comprehensive description of our supervised machine learning framework, significance tests, and information-theoretic approaches, please refer to the Information Quantification.

## Declarations

### Supplementary information

The supplementary information for this paper includes the appendix for the Information Quantification and extended data Figures.

### Funding

This research was funded by the Postdoctoral Junior Leader Fellowship Programme from La Caixa Banking Foundation (LCF/BQ/PI18/11630004).

### Competing interests

The authors declare no competing interests. The funders had no role in the design of the study; in the collection, analyses, or interpretation of data; in the writing of the manuscript, or in the decision to publish the results.

### Ethics approval

Not applicable

### Consent to participate

Not applicable

### Consent for publication

Not applicable

### Data availability

Not applicable

### Code availability

Code supporting the findings of this paper are available on github (61, 65).

### Authors’ contributions

F.R. and B.S. conceived the research. B.S. secured funding. F.R. and B.S. developed the methodology. F.R. designed the simulations and carried out data curation, data analysis, and data visualization. F.R. developed the Python package for information quantification. F.R. wrote the original draft. F.R. and B.S. wrote and edited the manuscript. B.S. supervised the research.

## Acknowledgment

We thank Francesco Damiani for comments on this manuscript.

## Information Quantification

Neural activity can be spontaneous or stimulus-driven. In a simplified hierarchical processing model, neurons respond to external stimuli by generating firing patterns. These patterns then influence the activity of downstream neurons, creating a cascade of information flow (68, 69). Postsynaptic neural spikes encode information about presynaptic firing, which in turn reflects the original external stimuli. We adopt the perspective that information, from a neuron’s viewpoint, is *“a difference that makes a difference”* (64). To quantify this information in neural responses, we introduce two novel measures: *information detection* and *information differentiation*.

### Information detection

Information detection assesses whether stimulus-induced neural firing significantly differs from spontaneous activity. It determines if neural responses can be statistically linked to stimulus presentation. While crucial for perception, information detection alone doesn’t guarantee rich stimulus encoding. For instance, if responses to two distinct stimuli both differ from spontaneous activity but not from each other, stimulus discrimination is limited. This scenario indicates attenuated, rather than absent, information content, as it still distinguishes stimulus-driven from spontaneous activity.

### Information differentiation

Information differentiation evaluates whether neural responses to different stimuli are statistically distinguishable. It measures the ability to attribute neural firing to specific stimuli within a given set. The degree of differentiation may vary with the introduction of new stimuli. Together, information detection and differentiation determine the overall information content in neural responses.

High levels of both enable accurate stimulus feature decoding by an ideal observer with prior stimulus knowledge. This framework applies to both spike-based and rate-based theories, encompassing signal and noise correlation coding schemes.

### Information propagation

Information flow between neural groups in a hierarchical processing chain can be assessed by measuring information content under different conditions. Consider a scenario where information content in a presynaptic group remains constant across two brain states. If the postsynaptic group maintains information detection but loses differentiation in one state, it indicates attenuated information content and reduced information flow from presynaptic to postsynaptic neurons in that state. Both detection and differentiation can decrease (or increase) simultaneously during state transitions.

To quantify information detection and differentiation, we employ machine learning algorithms. Information detection is measured by an algorithm’s ability to distinguish stimulus-induced firing from spontaneous activity. Information differentiation is assessed by the algorithm’s capacity to discriminate between responses to different stimuli. For researchers interested in replicating or extending our analysis, we have made our custom Python module for information quantification, *iQuanta*, publicly available on GitHub (65).

### Supervised Machine Learning framework

In our supervised approach, we employed the Generalized Linear Model (GLM) to quantify neural information content. This method involves three key steps: model selection, parameter estimation, and prediction (70). The assumption of independent observations in GLM was met through independent simulations across trials.

For information detection, we employed a GLM with a Bernoulli distribution for evoked firing responses at stimulus offset to a specific stimulus intensity and spontaneous firing activities in each brain state. We fitted a binomial logistic classification model to training sets, optimizing parameters via Newton-Raphson to maximize log likelihood (71, 72). Performance was assessed using stratified 10-fold cross-validation, with classification accuracy averaged across folds (67). Higher accuracy indicates greater information detection (see Extended Data Fig. 9a(i)).

To quantify information differentiation, we applied a GLM separately for each brain state. We used a multinomial distribution for evoked firing responses at stimulus offset across the battery of stimulus intensities and optimized parameters via the Newton-Raphson method. Model performance was assessed using stratified 10-fold cross-validation, with classification accuracy scores averaged across folds. Higher accuracy indicates greater information differentiation (see Extended Data Fig. 9a(ii)).

### Significance Test

Moving beyond machine learning approaches, we performed statistical significance tests on the firing rate responses at stimulus offset to quantify changes in information content.

To quantify the difference between evoked responses and spontaneous prestimulus activities, we applied Student’s t-test for each stimulus intensity in each brain state. With 500 trials satisfying the Central Limit Theorem, we calculated independent t-test statistics (t-values). Statistical significance was set at p < 0.05. The *t*-value served as a measure of the distinction between evoked and spontaneous activities, with higher *t*-values indicating greater information detection (see Extended Data Fig. 9b(i)).

To evaluate the distinctiveness of evoked responses across stimulus intensities, we conducted one-way ANOVA separately for each brain state. We calculated the F-ratio as a measure of response differentiation. Higher F-ratios indicate greater information differentiation between stimuli (see Extended Data Fig. 9b(ii)).

### Information Theory

We also employed information theory to assess the informativeness of evoked firing responses at stimulus offset about stimulus intensities. Mutual information quantified the relationship between firing rate distributions and stimulus intensities (69). Greater separation among stimulus-evoked response distributions yields higher mutual information, potentially indicating increased information differentiation (see Extended Data Fig. 9c). For numerical computation, firing rates at stimulus offset were discretized into 0.1 Hz bins to calculate probabilities.

## Extended Data Figures

**Extended Data Figure 1.**
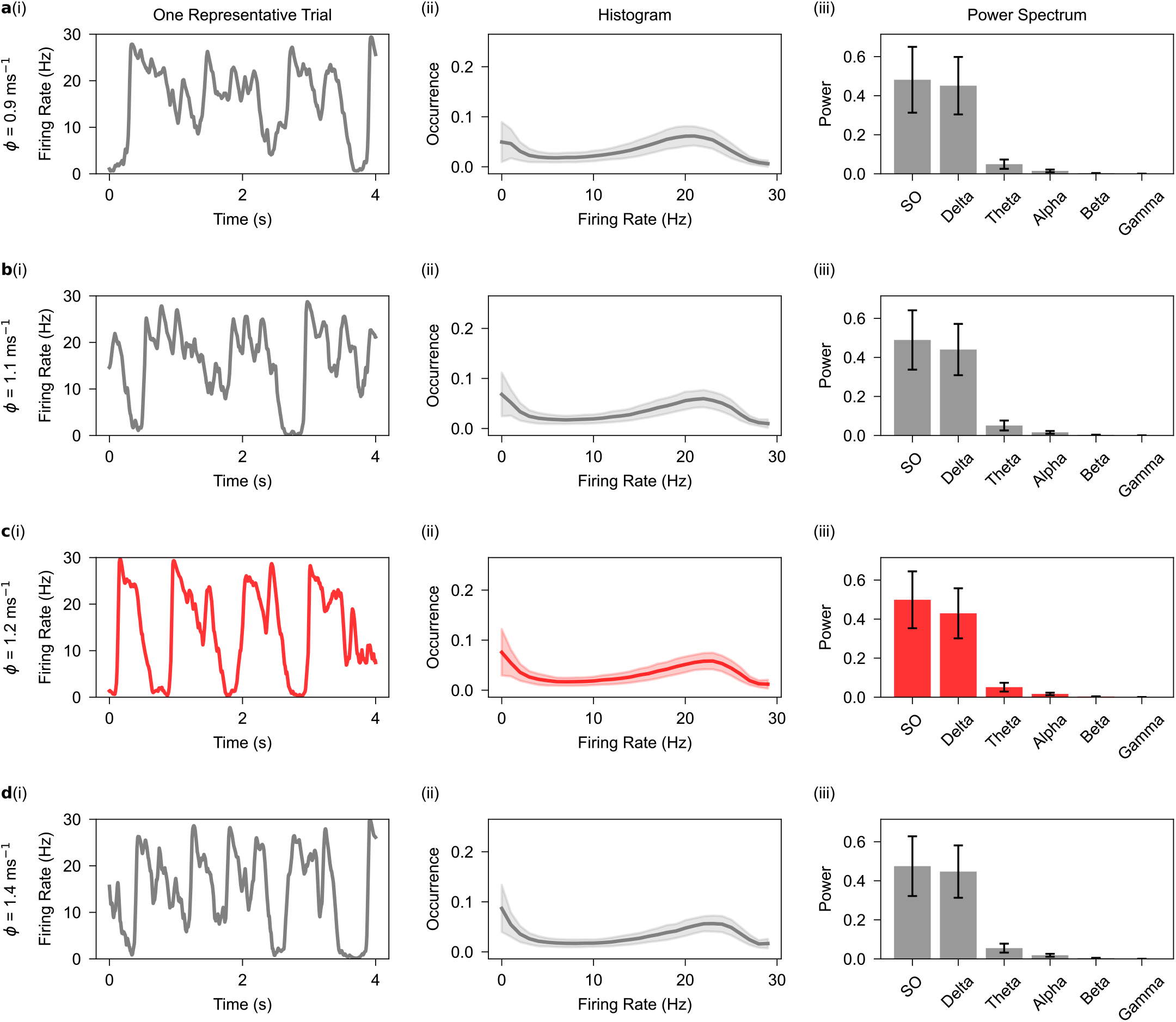
Dynamical features of spontaneous firing activity in the one-cortical-column model are robust to the changes in the standard deviation of the noise, *ϕ*. **a**, Spontaneous firing rate signal for a representative trial (i), the distribution of firing rate signals (ii), and the power spectrum of signals (iii) when *ϕ* = 0.9 ms^−1^. **b, c**, and **d**, As in **a**, but for when *ϕ* increases. Panel **c** here is as Fig. 1**c**. Note that *ϕ* = 1.2 ms^−1^ is used as the value of the standard deviation of the noise in this computational study. Shaded area and Error bar correspond to standard deviation over 500 trials.

**Extended Data Figure 2.**
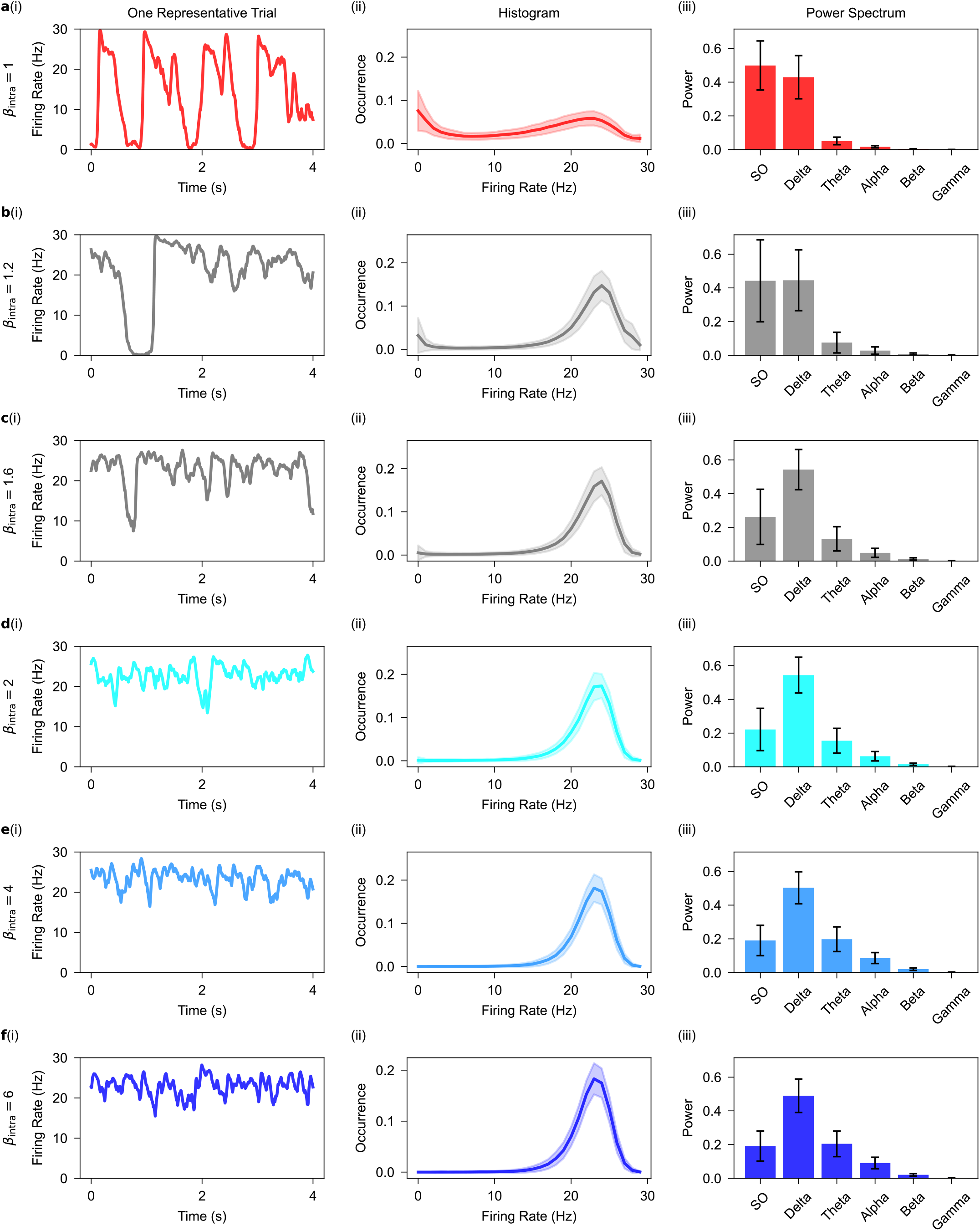
Dynamical features of spontaneous firing activity in the one-cortical-column model changes with increasing intra-synaptic upscaling, *β*_intra_. **a**, Spontaneous firing rate signal for a representative trial (i), the distribution of firing rate signals (ii), and the power spectrum of signals (iii) when there is no intra-synaptic upscaling (*β*_intra_ = 1). Panel **a** here is as Fig. 1**c. b, c, d, e**, and **f**, As in **a**, but for when intra-synaptic upscaling (*β*_intra_) increases. Panel **d** here is as Fig. 1**d**. Shaded area and Error bar correspond to standard deviation over 500 trials.

**Extended Data Figure 3.**
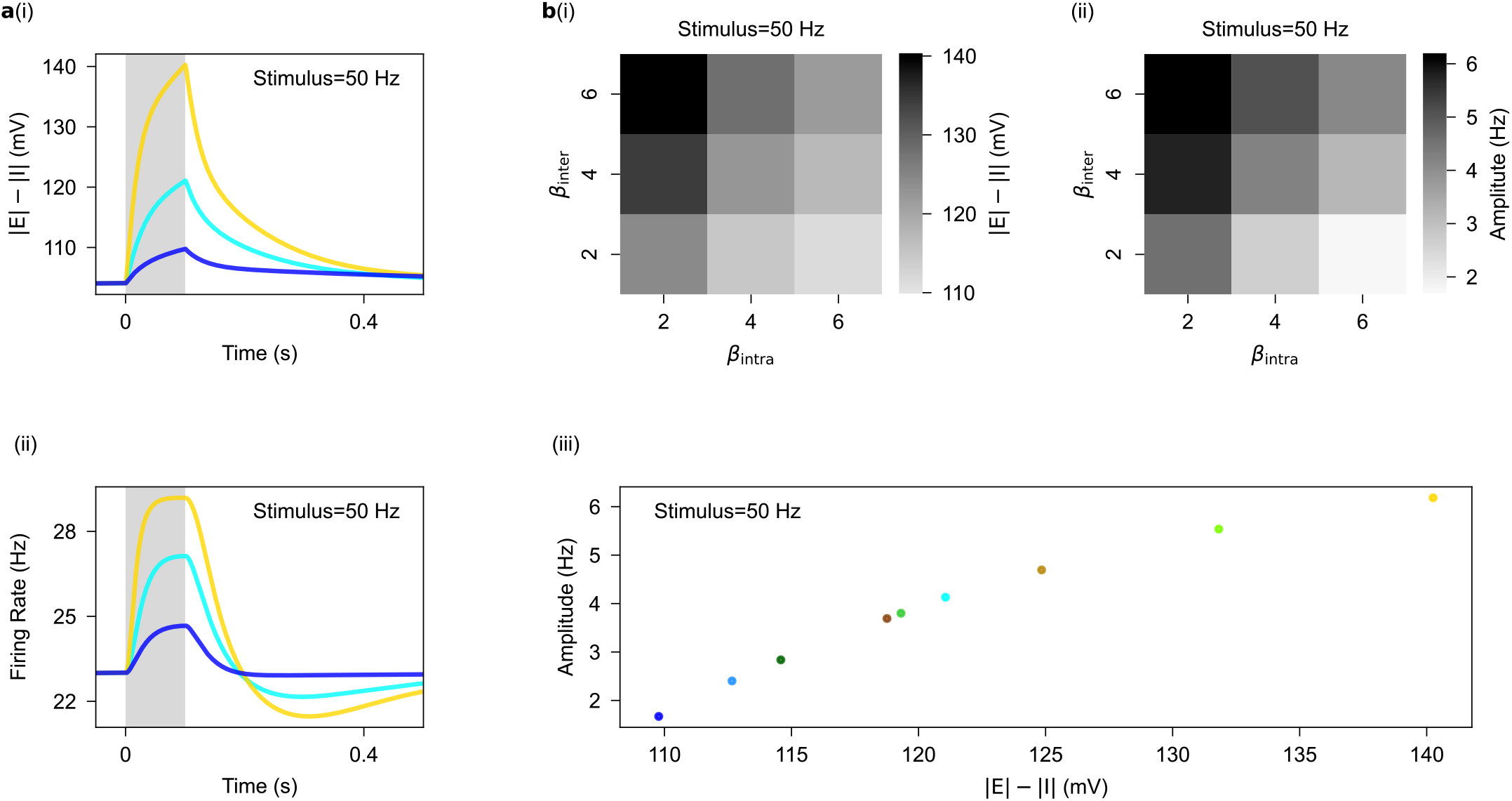
Effect of intra- and inter-synaptic upscaling on the net evoked synaptic currents in the one-cortical-column model. **a**, The net evoked synaptic current (i), quantified as |E| −|I|, decreases with increasing *β*_intra_ (from ligh to dark blue) as opposed to when *β*_inter_ increases (from ligh blue to light yellow) in wakefulness. Changes in the time trace of net evoked synaptic currrent determine changes in the time trace of evoked firing responses (ii). This holds true for other values of stimulus intensity (not shown here). Note that the net synaptic current remains constant before stimulus onset across various synaptic upscaling scenarios, illustrating that synaptic upscaling is implemented in a balanced configuration without causing predominant excitation or inhibition. Shaded area corresponds to the stimulus duration. **b**, Effects of *β*_intra_ and *β*_inter_ on the net evoked synaptic currents explain the pulling and driving effects associated with the intra- and inter-synaptic upscalings in wakefulness. Intra-synaptic upscaling decreases the net evoked synaptic current (i) that results in the decreased evoked responses (ii). Conversely, inter-synaptic upscaling increases the net evoked synaptic current (i) that results in the increases evoked responses (ii). Changes in the net evoked synaptic currrent determine changes in the amplitude of evoked firing responses (iii). Note that analysis in **b**(i) are carried out on the data points at stimulus offset.

**Extended Data Figure 4.**
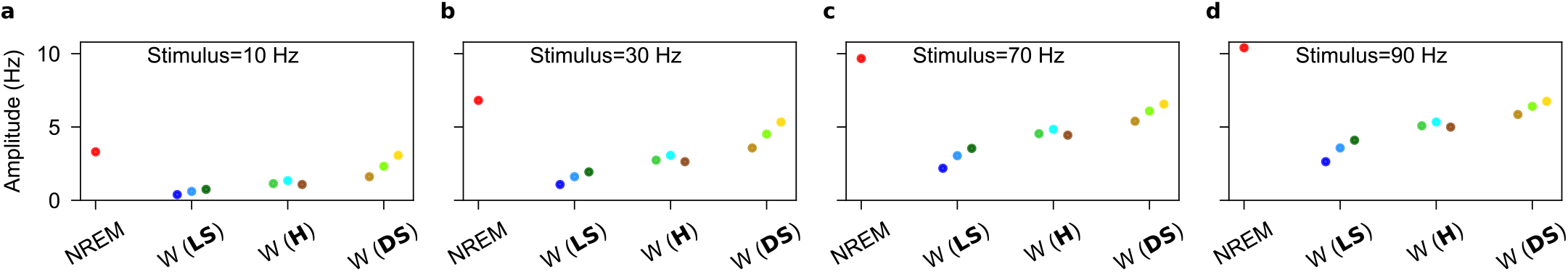
The amplitude of evoked firing responses in the one-cortical-column model. **a**, The amplitude of evoked firing responses increases with increasing values of synaptic upscaling ratio, *β*_inter_*/β*_intra_, during wakefulness when the stimulus intensity is 10 Hz (**a**), 30 Hz (**b**), 70 Hz (**c**) and 90 Hz (**d**). Note that the overall enhancement of the amplitude of evoked responses as the stimulus intensity increases from **a** to **d**.

**Extended Data Figure 5.**
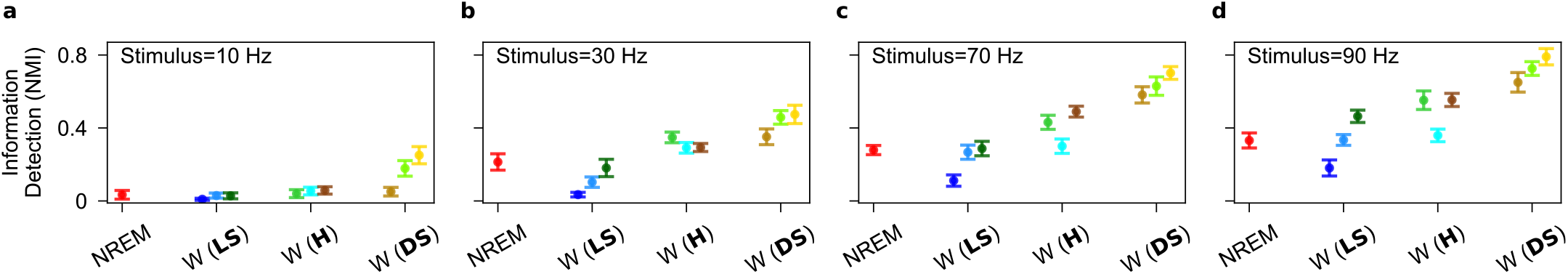
Information detection in the one-cortical-column model. **a**, Information detection increases with increasing values of synaptic upscaling ratio, *β*_inter_*/β*_intra_, during wakefulness when the stimulus intesity is 10 Hz (**a**), 30 Hz (**b**), 70 Hz (**c**) and 90 Hz (**d**). Synaptic upscaling during wakefulness does not enhance information detection during wakefulness across stimuli compared to those in NREM sleep unless it occurs in DS upscaling. Note that the overall enhancement of information detection as the stimulus intesity increases from **a** to **d**. Error bar corresponds to 95% confidence interval over 10 performance estimate of the K-means clustering algorithms.

**Extended Data Figure 6.**
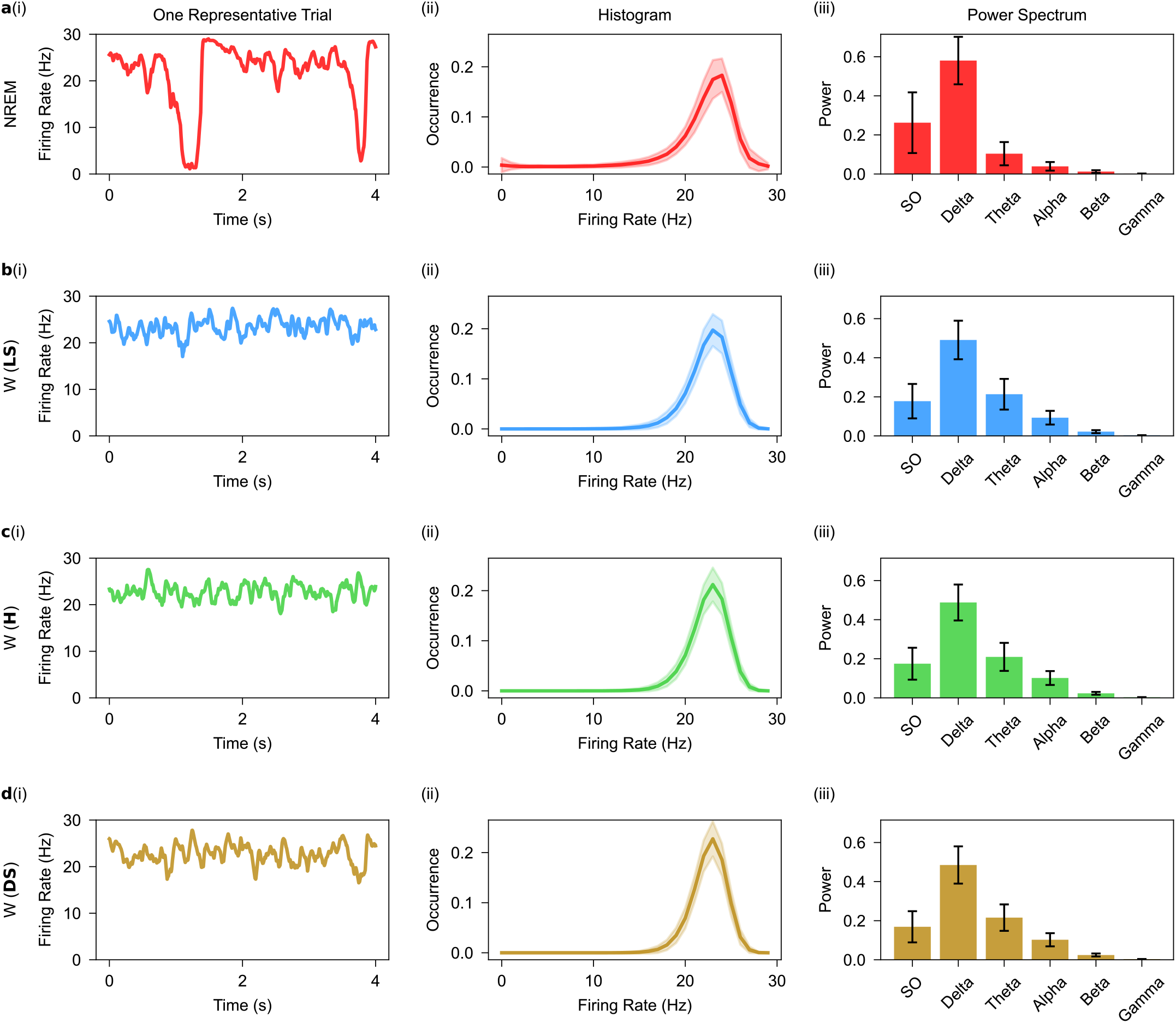
Dynamical features of spontaneous firing activities in the two-cortical-column model. **a**, Spontaneous firing rate signal for a representative trial (i), the distribution of firing rate signals (ii), and the power spectrum of signals (iii) when there is no synaptic upscaling (*β*_intra_ = 1, *β*_inter_ = 1). **b, c**, and **d**, As in **a**, but for when synaptic upscaling is local-selective (LS: *β*_intra_ = 4, *β*_inter_ = 2), homogeneous (H: *β*_intra_ = 4, *β*_inter_ = 4), and distance-selective upscaling (DS: *β*_intra_ = 4, *β*_inter_ = 6), respectively. The dynamical features of spontaneous firing activity in the two-cortical-column model shift from NREM sleep to wakefulness for all synaptic upscaling combinations (not shown here). Shaded area and Error bar correspond to standard deviation over 500 trials.

**Extended Data Figure 7.**
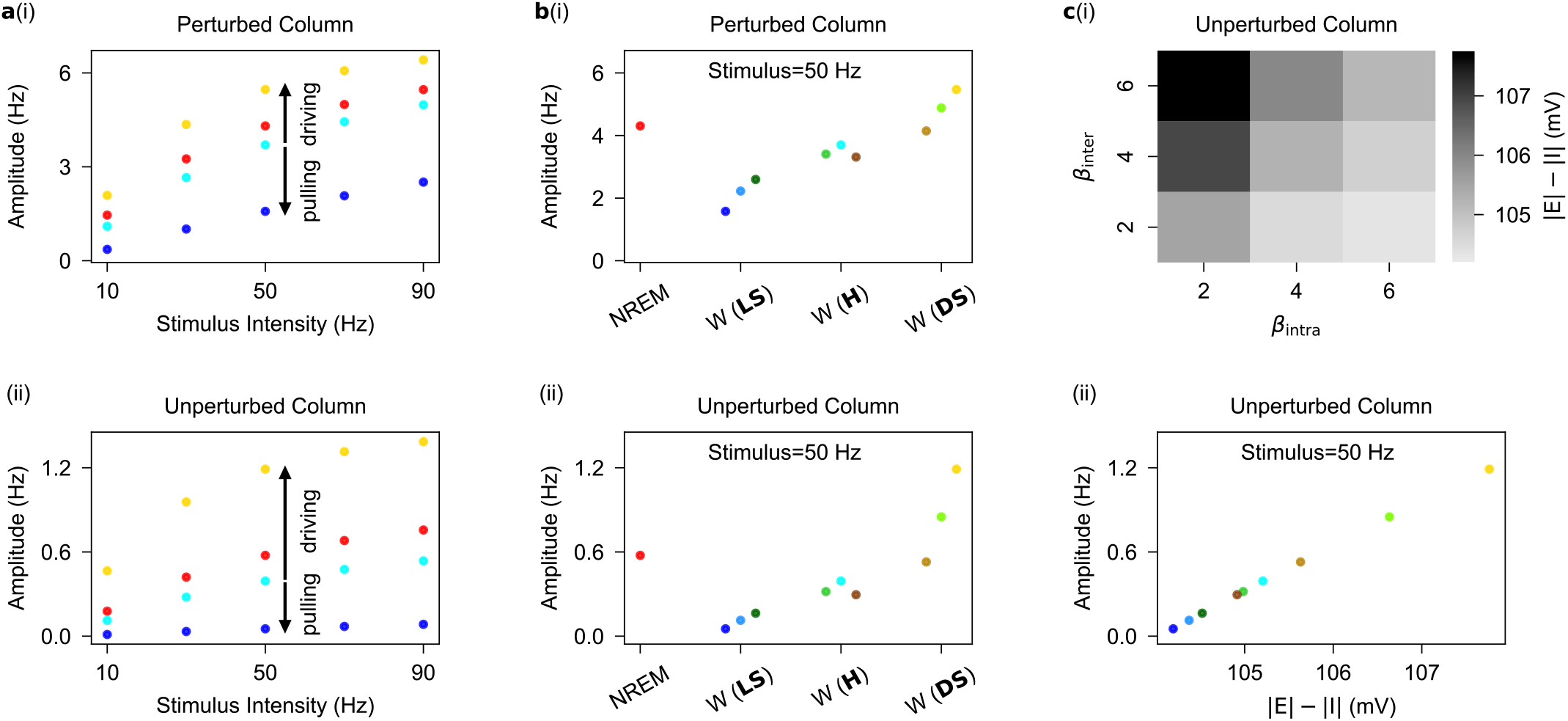
Effect of intra- and inter-synaptic upscaling on the response of two-cortical-column model to stimuli. **a**, Increasing intra-synaptic upscaling while inter-synaptic upscaling is constant from light to dark blue (from *β*_intra_ = 2, *β*_inter_ = 2 to *β*_intra_ = 6, *β*_inter_ = 2) during wakefulness produces a pulling effect on the amplitude of evoked firing responses in the perturbed (i) and unperturbed cortical column (ii). Conversely, increasing inter-synaptic upscaling while intra-synaptic upscaling is constant from light blue to light yellow (from *β*_intra_ = 2, *β*_inter_ = 2 to *β*_intra_ = 2, *β*_inter_ = 6) during wakefulness produces a driving effect on the amplitude of evoked firing responses in the perturbed (i) and unperturbed cortical column (ii). **b**, The amplitude of evoked firing responses increases as the synaptic upscaling transitions from local-selective (LS) to distance-selective (DS) upscaling during wakefulness in the perturbed (i) and unperturbed cortical column (ii). Note that this holds true for other values of stimulus intensity (not shown here). The amplitude of evoked responses to stimuli in the perturbed (i) and unperturbed cortical column (ii) during wakefulness enhances as synaptic upscaling transition from local-selective (LS) towards distance-selective (DS) upscaling. **c**, Inter-synaptic upscaling increases the net evoked synaptic current, as opposed to when intra-synaptic upscaling increases during wakefulness in the unperturbed cortical column (i). Changes in the net evoked synaptic currrent determine changes in the amplitude of evoked firing responses (ii).

**Extended Data Figure 8.**
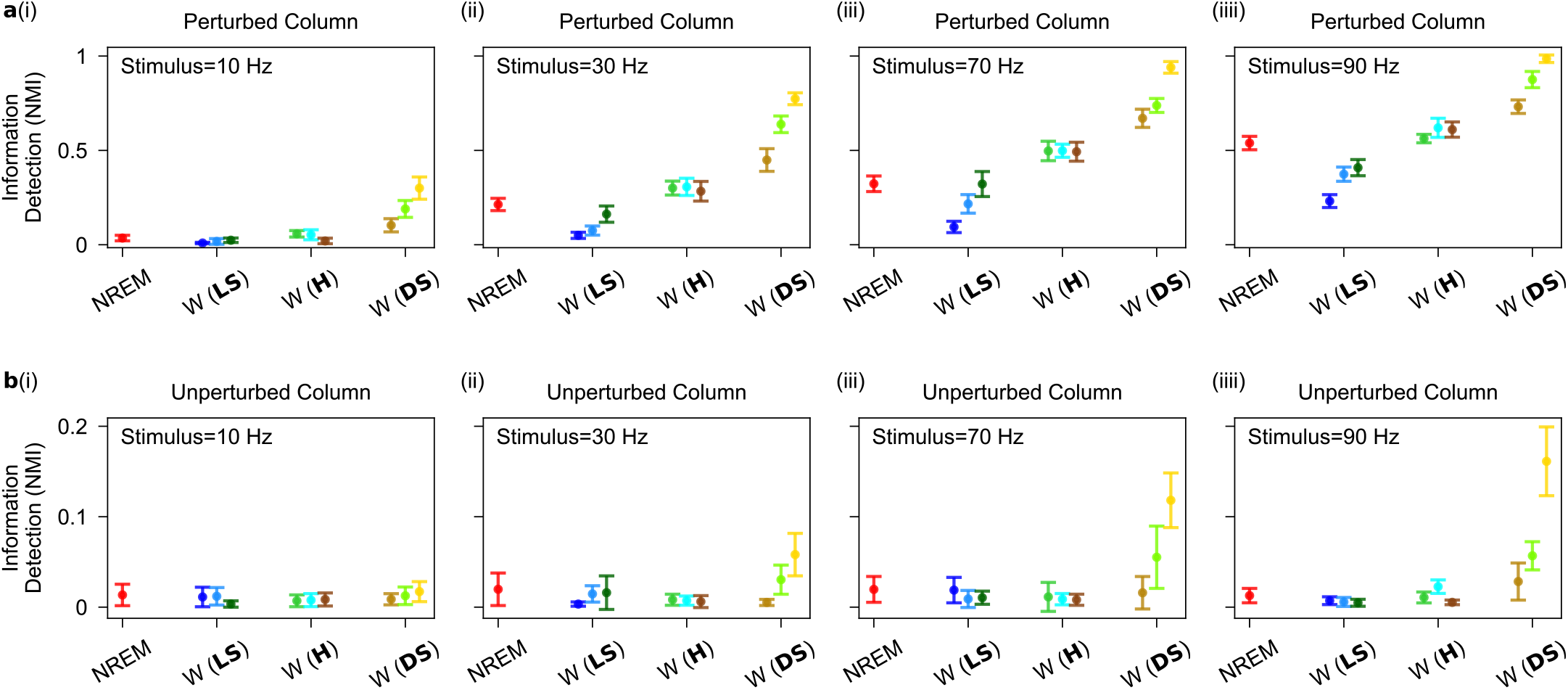
Information detection in the two-cortical-column model. **a**, Information detection increases with increasing values of synaptic upscaling ratio, *β*_inter_*/β*_intra_, during wakefulness when the stimulus intesity is 10 Hz (i), 30 Hz (ii), 70 Hz (iii) and 90 Hz (iiii). **b**, As in **a**, but for the unperturbed cortical column. Error bar corresponds to 95% confidence interval over 10 performance estimate of the K-means clustering algorithms.

**Extended Data Figure 9.**
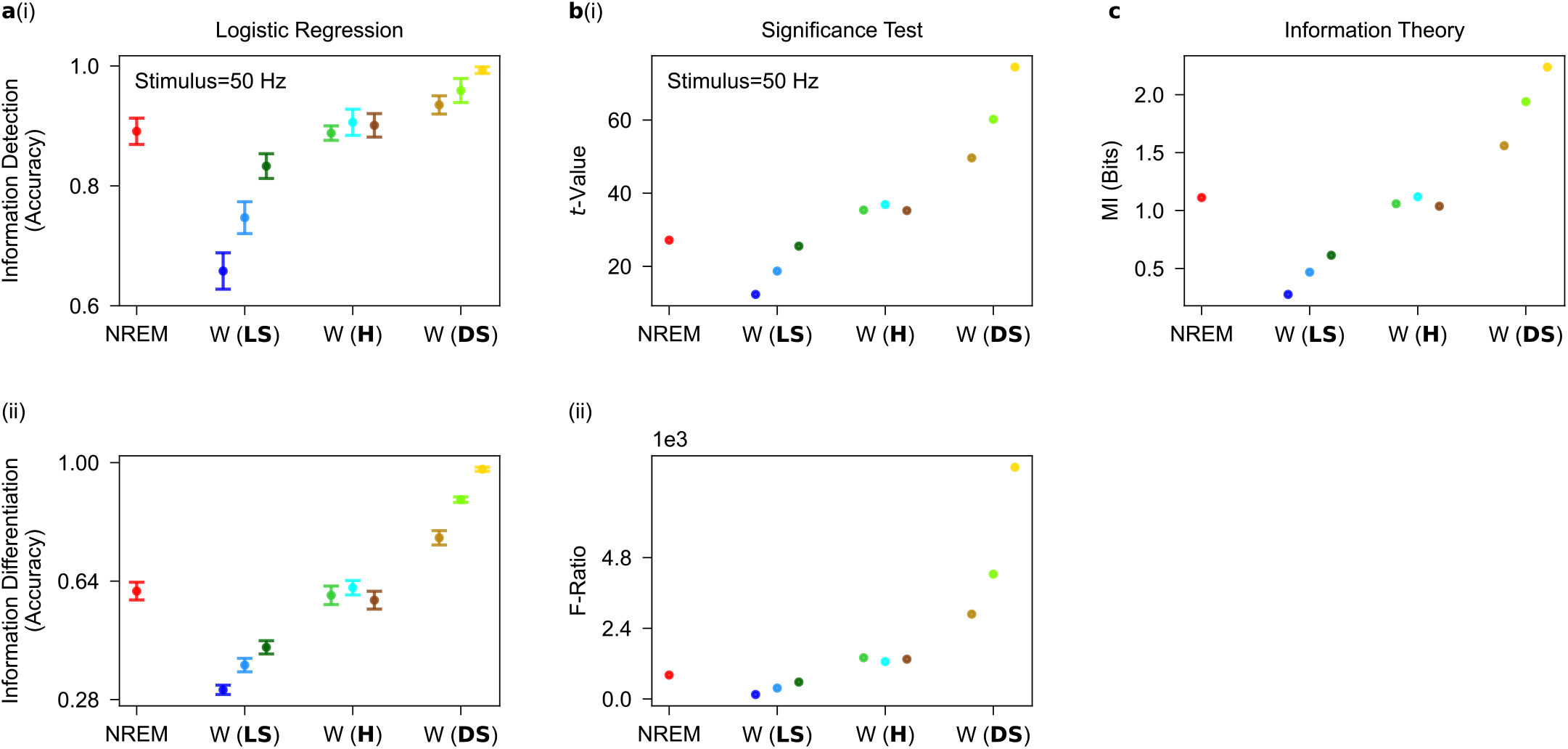
Robustness of the computational results. Figures pertain to analysis of evoked firing responses in the perturbed cortical column in the two-cortical-column model. **a**, Information detection for when stimulus intensity is 50Hz (i) and information differentiation (ii) when logistic classification algorithms (see Information Quantification) are employed. Error bar corresponds to 95% confidence interval over 10 performance estimate of the logistic classification algorithms. Logistic classification algorithms qualitatively replicate the results obtained using K-means clustering algorithms in Fig. 4**b. b**, Implementing significance tests (see Information Quantification) such as student t-test (i) and analysis of variance (ii) qualitatively replicate the results obtain by machine learning techniques pertaining to information detection and information differentiation. **c**, Implementing information theory (see Information Quantification) manifests that the mutual information between the distribution of evoked responses at stimulus offset and the distribution of stimuli increases as synaptic upscaling transitions from local-selective (LS) to distance-selective (DS) upscaling during wakefulness.

## Notes

### Competing Interest Statement

The authors have declared no competing interest.

### Summary of Updates

This revised manuscript features a restructured methodology, enhancing clarity and focus. The original method section is now divided into a concise Method section and a detailed Appendix. The Method section provides a streamlined overview of our approach, maintaining all essential elements while improving readability. Key improvements include a brief introduction to our framework for quantifying information content in the main Method section. This overview outlines the core principles and basic application of the framework, providing readers with a clear understanding of our analytical approach. The Appendix now contains a comprehensive description of the framework for quantifying information content. This detailed exposition is tailored for researchers interested in contributing to or expanding upon this field. It includes in-depth explanations of the mathematical foundations, step-by-step implementation guidelines, and discussions on potential extensions and applications of the framework. By separating the concise overview from the detailed exposition, we cater to different reader needs. The main Method section serves those seeking a general understanding of our approach, while the Appendix provides the depth required for researchers looking to engage more deeply with the framework. This restructuring not only enhances the accessibility of our core methodology but also provides a valuable resource for future research in information content quantification. The detailed Appendix serves as a foundation for further development and application of the framework in various contexts.

